# Dispersal evolution can only rescue a limited set of species from climate change

**DOI:** 10.1101/2024.12.05.626982

**Authors:** Peter Kamal, Patrick L. Thompson, Natalie Lewis, Emanuel A. Fronhofer

**Affiliations:** ISEM, University of Montpellier, CNRS, IRD, EPHE, 34095 Montpellier, France; Fisheries and Oceans Canada, Pacific Science Enterprise Centre, West Vancouver, BC V7V 1H2, Canada; Department of Ecology & Evolution, University of Lausanne, Biophore, 1015 Lausanne, Switzerland

## Abstract

Anthropogenic climate change threatens biodiversity on Earth. In response, species can adapt evolutionarily to changing environments or shift their ranges via dispersal. However, dispersal itself can evolve on ecological timescales. We explore theoretically how dispersal evolution modulates the response of metacommunities to climate change. We find that this response depends on the environmental conditions prior to climate change. Variable environments harbour few, dispersive species that are likely to survive climate change by shifting their ranges with their evolved dispersal abilities. Stable environments house many, less dispersive species. Their survival during climate change is less likely, as they can evolve robust, low dispersal traits that prevent range shifts. We identify a limited set of scenarios in which contemporary dispersal evolution can rescue species from climate change, highlighting the importance of species’ evolutionary histories and evolutionary rates, as determined by their genotype-phenotype maps, for their responses to rapid environmental change.

## Introduction

One of the main drivers of current biodiversity loss is the rise in temperatures due to climate change [1, 2]. Understanding which species survive under increasing temperatures, and why, is crucial for forecasting and potentially preventing further species extinctions [3, 4]. Generally, organisms can survive changing conditions (i) via local adaptation, that is, evolutionarily adapt to new conditions by shifting their niches to track environmental change [5, 6] or (ii) via dispersal, that is, species shift or expand their range and disperse to habitats they are already adapted to, thereby tracking conditions in space [7, 8]. However, any species’ response to environmental change is heavily modulated by biotic interactions such as competition [9, 10]: If suitable habitat is already occupied, dispersing into this habitat may not be an option.

Theoretical work on the evolutionary ecology of metacommunities under climate change has generated one relatively consistent prediction: Dispersal and local adaptation can be antagonistic for the maintenance of biodiversity [11–13]. In particular, the interplay of these two processes can produce monopolization effects in space: Early-arriving species may adapt to the local conditions and then prevent later-arriving species from colonizing [14–16]. As a consequence, these evolution-mediated priority effects can contribute to biodiversity loss [12, 17].

This theoretical work has generally assumed invariant dispersal rates. However, dispersal is genetically determined [18, 19] and can evolve on ecological timescales, particularly during range expansions [20, 21]. Single-species models have presented proof-of-concept that rapid dispersal evolution can rescue populations during climate change [22, 23]. Multi-species models have, so far, only studied dispersal evolution during metacommunity assembly but show important interaction effects with local adaptation and the abiotic and biotic environment [24, 25]. To better forecast biodiversity loss, it is crucial that we understand how these interactions impact the potential of dispersal evolution to rescue species during climate change.

Dispersal evolution might rescue species during climate change by allowing them to perform range shifts to more suitable habitats that would have been impossible without rapid evolutionary change [22, 23]. This rescue mechanism would reinforce itself because of spatial sorting of individuals with strong dispersal abilities at the range front, speeding up the pace of evolutionary change [20, 21]. Theoretically, the pace of dispersal evolution could therefore at least track the pace of climate change [22], and such tendencies have been shown in some natural populations [26, 27]. This potential of dispersal evolution to match the speed of climate change is particularly important given that climate change is generally predicted to outpace the rates of climatic niche evolution in many species [28–30], leading to possibly severe losses in global biodiversity [31, 32].

Here, we use a stochastic, spatially explicit, individual-based model to explore how the joint evolution of local adaptation to an environmental gradient and dispersal modulates the response of a metacommunity to climate change. We ask the following questions: (1) How does dispersal evolution shape community assembly; and by extension, how do the drivers of dispersal evolution — particularly environmental stochasticity — shape community assembly? (2) How do species respond to climate change when both dispersal and local adaptation can evolve? (3) Can contemporary dispersal evolution rescue species from climate change, and if so, under which conditions? The complexity of the underlying mechanisms gives rise to a suite of possible outcomes: The most positive outcome for biodiversity maintenance would be that, as in the single-species case, dispersal evolution could allow species to be rescued from climate change [22, 23] despite — or even fuelled by — competitive pressure and a lack of evolutionary potential for local adaptation. Conversely, it is possible that local adaptation and competitive pressures during range expansions constrain the ability of dispersal evolution to impact biodiversity maintenance in any meaningful way, similar to the antagonistic interaction between local adaptation and dispersal shown by Thompson & Fronhofer [12]. By answering these questions, we refine our mechanistic understanding of the complex processes governing biodiversity maintenance — which is urgently needed for both fundamental and applied purposes [4].

## Materials & Methods

We use a stochastic, spatially explicit, individual-based metacommunity model with an underlying genetic algorithm assuming non-overlapping, discrete generations. The model was first presented in Thompson & Fronhofer [12] and featured only local adaptation as an evolving trait. Here, we add dispersal as a second evolving trait. Figure 1 sketches the main elements of the model.

**Figure 1:**
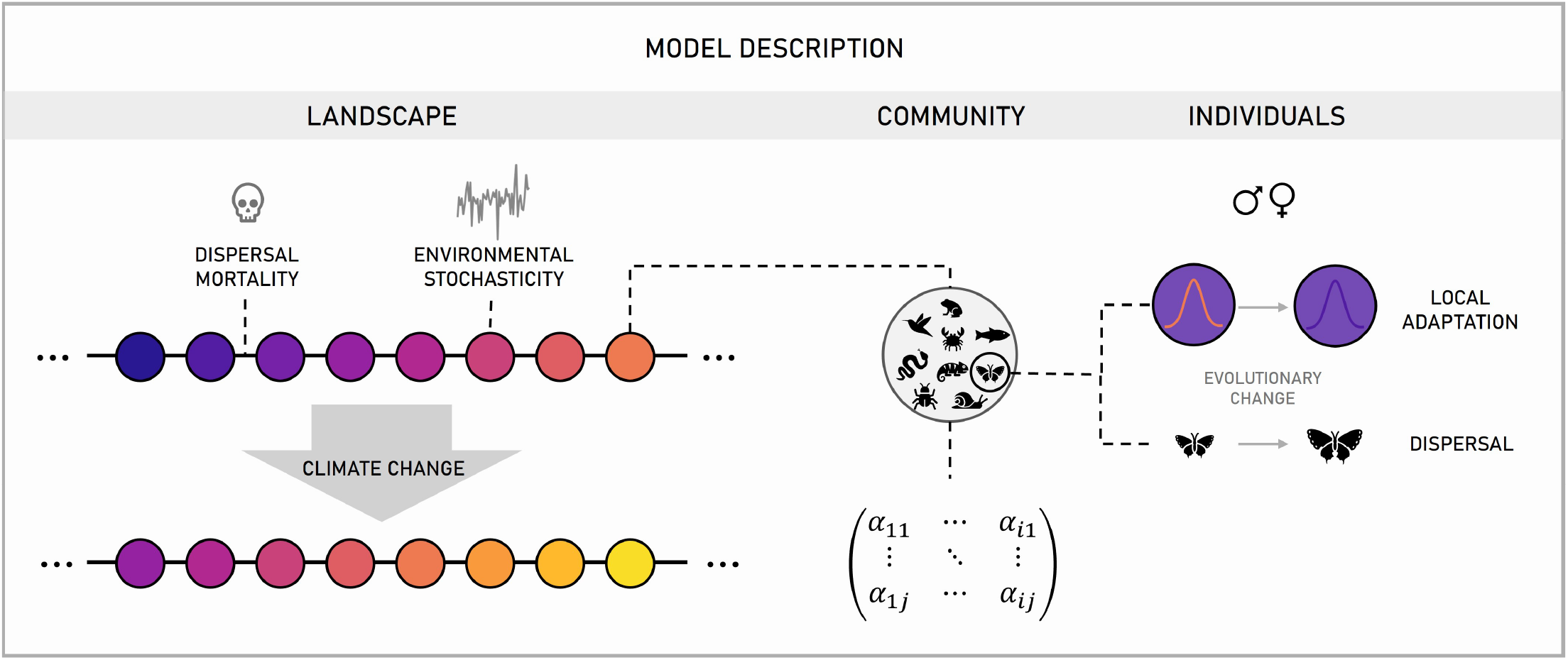
Graphical sketch of the model. We model a metacommunity across a temperature gradient. Each local community interacts according to a Lotka-Volterra-type competition matrix. Individuals reproduce sexually. Each individual has two traits that are under selection: local adaptation to the environment, and dispersal. Dispersal mortality and environmental stochasticity select against and for dispersal, respectively. After the metacommunity has undergone eco-evolutionary assembly, the temperatures increase across the landscape to simulate climate change. We depict here only a part of the landscape for graphical simplicity. The full landscape is a ring, along which temperatures first increase and then decrease (see Figure S1).

### Time and Space

In the model, we track 80 initial species through time and space. The world is a ring of 30 patches (so to avoid edge effects) with a gradient in local conditions that could, for example, represent temperature but also any other environmental factor. Here, we chose values that increase in integer steps from 0 to 15 and then decrease again. The ring shape of the gradient effectively creates geographic poles that can be approached from both sides (see Figure S1). Thompson & Fronhofer [12] showed that the general dynamics of our model also hold for a more complex landscape structure. One replicate simulation runs for a total of 15,000 generations. During the first 10,000 generations, temperatures are held constant to allow the system to obtain quasi-equilibrium. Climate change starts at generation 10,000: From then on, temperatures increase uniformly and gradually at a rateδ_*envchange*_ across the landscape until the end of the simulation.

### Species interactions

Species interact through competition. Each replicate run uses a unique, constant, and symmetric matrix of intra- and interspecific competition coefficients. These coefficients are drawn from lognormal distributions with variance 0.1. The distribution of intraspecific coefficients has a mean of 0.002, the distribution of interspecific coefficients has a mean of 0.001. This setting allows for coexistence on average, but not for every species pair. Each species *s* is initialised in every patch at the beginning of the simulation at its single-species equilibrium density 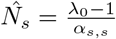 with *λ*_0_ = 2 the average fecundity of a female — a realistic value in natural populations [33] — and *α*_*s,s*_, the intraspecific competition coefficient.

### Genetic architecture

Individuals carry two diploid loci, one coding for emigration probability, one for local adaptation to the environment. We employ the simplest possible genotype-phenotype map: The phenotype of an individual is the arithmetic mean over its two alleles at each locus. Individuals of one species are initialised with the same local adaptation value, drawn randomly from a uniform distribution between 0 and 15 (the range of temperatures in the world before climate change). Dispersal is initialised at full standing genetic variation i.e., each allele of the first generation is assigned a dispersal probability randomly drawn from a uniform distribution between 0 and 1.

The values of both loci are unbounded on a genotypic level. Dispersal, being a probability, is bounded between 0 and 1 on a phenotypic level. We chose to not bound the dispersal genotype as initial tests with restrictive boundary conditions revealed that such bounds produce a bias in the optimal dispersal rates (Figure S2). We also tested a logit transformation of the dispersal trait, however as our evolutionarily stable dispersal rates are close to zero, this type of boundary condition effectively reduced the variation introduced by mutation to zero. Therefore, we chose to let the allelic values of the dispersal trait drift below zero. To reduce the drift, we use a simple annealing process, by which the mutation rate of dispersal (initial value = 0.001) linearly decreases to 0 between generations 2,500 and 5,000 [34]. Dispersal traits below zero are more robust to mutations [35] than traits at zero or above because changes to the genotype are not expressed in the phenotype and therefore not subject to selection. The annealing process ensures that the dispersal traits do not drift too far below zero which would make evolutionary rescue during climate change impossible. Between generation 5,000 and 10,000 (the start of climate change), selection acts only on the remaining variance in the populations. Once climate change starts, we allow for dispersal mutations at varying rates. We do not use a similar annealing process for local adaptation, as varying rates of local adaptation can produce different metacommunity dynamics even pre-climate change if environment is stochastically perturbed.

### Reproduction

Individuals reproduce sexually under random local mating. The reproductive output of a female follows a Beverton-Holt model [36]. To simulate environmental stochasticity, we draw a fecundity *λ*_*p*_(*t*) specific to each time step *t* and patch *p* from a lognormal distribution with mean fecundity *λ*_0_ = 2 and standard deviation *σ*. This equates to a patch-level environmental white noise. We then draw the number of offspring of a female from a Poisson distribution with the parameter *λ*_*p*_(*t*) and correct for intra- and interspecific competition and local adaptation to the environment so that the expected value of the draw for a female *f* of species *s* follows:

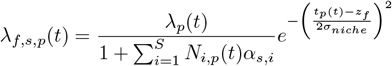

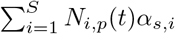 is the amount of local competition faced i.e., the product of population densities *N*_*i,p*_ for all species *i* ∈ [1, …, *S*] in patch *p* at time *t*, and *α*_*s,i*_ the relevant competition coefficients. *t*_*p*_(*t*) − *z*_*f*_ describes the environmental mismatch between *t*_*p*_(*t*), the temperature experienced in the patch by the individual at time *t*, and *z*_*f*_, the thermal phenotype of the individual. *σ*_*niche*_ describes the niche width, is set to 1 and held constant. This implies a reduction of ca. 22% in fitness for an optimally adapted individual that disperses to its neighbouring patch.

Offspring inherit one randomly drawn allele from each parent at each locus. During inheritance, mutations can occur. Mutations add a value drawn from a normal distribution with mean 0 and standard deviation 0.06 to the allelic value of the parent. The rate of mutation is set with a parameter specific to each trait. The mutation rates in our model are generally above those found in natural populations, but the standard deviation is set so that the evolutionary change in our traits is between 0 and 0.025 haldanes [12], which mirrors rates of contemporary microevolution in natural populations [37].

### Dispersal

Dispersal is natal and costly [38]. Individuals can only disperse to their two neighbouring patches, the choice of which is random. Dispersal in our model represents specifically emigration propensity i.e., the probability that an individual leaves its natal patch. It is consensus that dispersal is comprised of three stages: Emigration, transfer, and settlement [39, 40]. We explicitly model the emigration stage and implicitly cover the other two stages: transfer and its cost is proxied by the dispersal mortality *µ*_0_, which can also be interpreted as a crude proxy of habitat fragmentation [41]. Successful settlement and subsequent survival of offspring depend on other model properties: local adaptation, local competition, and the ability to find mates.

In our model, dispersal evolution is mainly driven by dispersal mortality *µ*_0_ — which selects against dispersal [38] — and by environmental stochasticity *σ* - which selects for increased dispersal [19, 39]. We do not vary dispersal mortality in the results presented in the main text, but the Supplementary Material contains sensitivity analyses in which this parameter is varied for all results.

### Outcome variables

We track the dynamics of the model on a metacommunity, species, population, and individual level. The most important outcome variables (per generation) for this paper are: (i) evolved emigration rate i.e., the number of dispersing individuals divided by the number of individuals in the metacommunity; (ii) mean local adaptation defined as 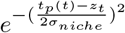 (see above for description), averaged over all females; (iii) mean *α*-diversity i.e., the average number of species per patch; (iv) *γ*-diversity i.e., the total number of species in the metacommunity; (v) community similarity, constructed as the ratio of *γ*- and *α*-diversity; (vi) mean range size i.e., the average number of patches occupied by one species; (viii) turnover rate, calculated as the ratio of number of patches per generation that change occupancy status compared to the full number of patches.

### Alternative scenarios

In our main scenario, all traits can evolve during climate change. We ran control scenarios, in which (i) no traits, (ii) only local adaptation, or (iii) only dispersal can evolve. This allows us to disentangle the impact of each trait evolving on metacommunity dynamics during climate change. As some environments produce ongoing priority effects, we use the community assembly phase to filter out the effects of the environment. Figure S3 illustrates this approach. To understand the role of priority effects better, we also ran an alternative scenario for community assembly, in which environmental stochasticity *σ* is replaced with random patch extinctions occurring at a rate *ϵ*.

### Parameters and simulations

Apart from the fixed parameters described in the text, Table S1 summarises the parameters we varied and the range of values we explored. We performed 50 replicate simulations each of our 3060 main parameter sets, and 25 for the 7120 parameter sets with varying speeds of climate change.

## Results & Discussion

To understand a metacommunity’s response to climate change, we must first understand the processes and outcomes of community assembly, as it sets the scene for any adaptation to climate change. Therefore, we first analyse the metacommunity pre-climate change depending on different levels of environmental stochasticity. Then, we explore the response to climate change and the drivers of this response for two representative environmental scenarios: invariable (low environmental stochasticity) and variable (high environmental stochasticity). The Supplementary Material contains sensitivity analyses for alternative values of environmental stochasticity for all results.

### Environmental stochasticity shapes metacommunity assembly via dispersal evolution

During community assembly, total biodiversity in the metacommunity is lower in more variable environments (Figure 2A). The species that survive in such environments evolve higher dispersal abilities (Figure 2B) and occupy larger ranges (Figure 2C). The increased environmental stochasticity selects for higher dispersal rates, which is a classic result known as bet-hedging [19, 39]. The higher the environmental stochasticity and the evolved dispersal rates, the less species can coexist (Figure S4A) due to monopolization effects [25] and the more similar local communities are to each other (Figure S4B) due to mass effects [42, 43]. These effects therefore reduce total biodiversity in the metacommunity in an expectable manner [24, 25]. Changes in dispersal mortality do not qualitatively change the results (Figure S5).

**Figure 2:**
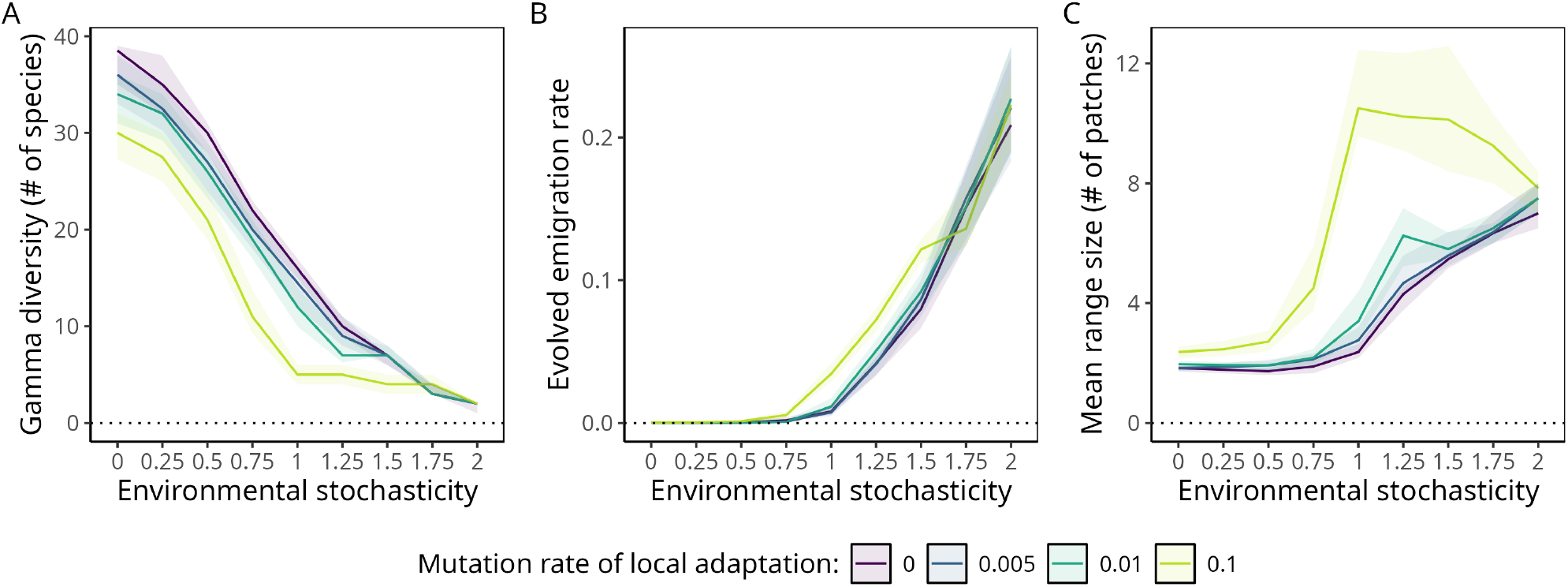
Community assembly dependent on environmental stochasticity. Lines represent medians and shaded areas interquartile ranges across 50 replicate simulations at the end of community assembly (t=10,000). Outcome variables are: (A) *γ*-diversity i.e., total number of species in the metacommunity; (B) the average emigration rate in the metacommunity; (C) the average range size of a species in number of patches. Dispersal mortality is set to an intermediate *µ*_0_ = 10%, and dispersal evolves with mutation rate of *m*_*d*_ = 0.001 that is annealed to 0 by t=5,000. For all other parameters see Table S1.

The loss of diversity at higher evolved dispersal rates is consistent with previous findings: Loreau *et al*. [44] find that at such high levels of dispersal, spatial insurance is not effective anymore in conserving regional biodiversity. Similarly, Amarasekare & Nisbet [45] find that dispersal rates above certain thresholds can inhibit coexistence between asymmetric competitors. In our model, priority effects might further compound these effects. We ran additional simulations, in which the environmental stochasticity was replaced by random patch extinctions, an extreme version of environmental noise [19]. The results are qualitatively similar, but a large amount of biodiversity is lost even when the rate of patch extinctions is very low (Figure S6). After such extinctions, the empty patches represent colonization opportunities, and therefore possible first-mover advantages. If dispersal rates are intermediate, local adaptation by the first movers might additionally reinforce these priority effects evolutionarily [15]. This reasoning might also explain the peak in range sizes at intermediate levels of stochasticity and high potential for local adaptation (Figure 2C). Our interpretation of the results matches natural and experimental observations from floodplains [46] and temporary ponds [47], typical instances of highly variable environments. In these systems, local communities were shown to become more similar across space and less diverse overall if given repeated and stochastic chances to recolonise habitat.

In concordance with previous research, our results demonstrate that environmental stochasticity is not only an important driver of dispersal evolution [39] but also of community assembly in heterogeneous environments [24, 25]. Different levels of stochasticity select for different dispersal rates and can subsequently produce a continuum of metacommunity dynamics from species sorting to mass effects [17, 42, 43]: Species sorting occurs in invariable environments that select strongly against dispersal and describes a case with many species occupying small, overlapping ranges. Mass effects occur in highly variable environments that select for dispersal, and describe a scenario with a few species that occupy large ranges and competitively exclude each other. These different dynamics in the community assembly process will prove crucial to understanding the metacommunity’s response to climate change.

### Climate change causes biodiversity loss and range expansions

During climate change, biodiversity in the metacommunity declines in both invariable and variable environments (Figure 3A). The only exception occurs when species have such high potential for local adaptation that they can perfectly track the changing conditions by rapidly adapting their niche (Figure 3A, green; Figure S7A). During climate change, dispersal rates tend to increase in invariable environments and stay mostly constant in variable environments (Figure 3B). We observe an increase in range size, particularly in invariable environments (Figure 3C), and these range expansions are directed towards the poles (Figure S7B). Changes to dispersal mortality or to the environment do not qualitatively change the outcomes, except for highly chaotic environments (Figure S8).

**Figure 3:**
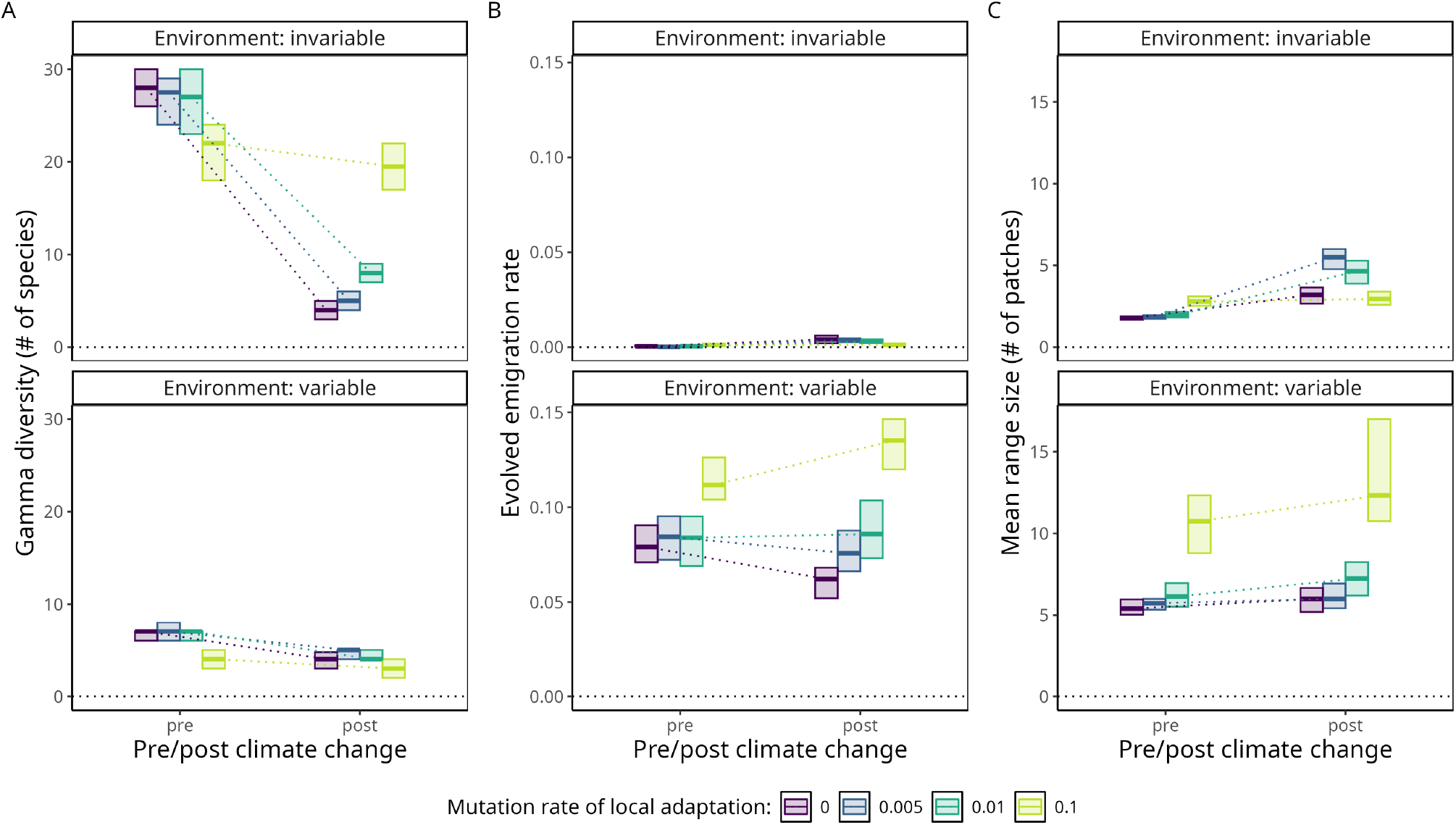
Response of the metacommunity to climate change. Thick lines represent medians and shaded boxes interquartile ranges across 50 replicate simulations pre (t=10,000) and post (t=15,000) climate change. Dispersal mortality is set to *µ*_0_ = 10% and the mutation rate of dispersal during climate change is set to *m*_*d*_ = 0.0005. In invariable environments, environmental stochasticity is set to *σ* = 0.5. In variable environments, environmental stochasticity is set to *σ* = 1.5. Outcome variables are: (A) *γ*-diversity i.e., total number of species in the metacommunity; (B) the average emigration rate in the metacommunity; (C) the average range size of a species in number of patches.

We identify two major determinants of species survival during climate change: First, species occupying ranges closer to the poles are more likely to go extinct during climate change (Figure 4A). Compared to more equatorial species, their capacity for poleward shifts is reduced. This effect is particularly pronounced in variable environments, where poleward range contractions reduce the capacity for bet-hedging. This result is rather unsurprising and matches a widely accepted finding from the literature [26].

**Figure 4:**
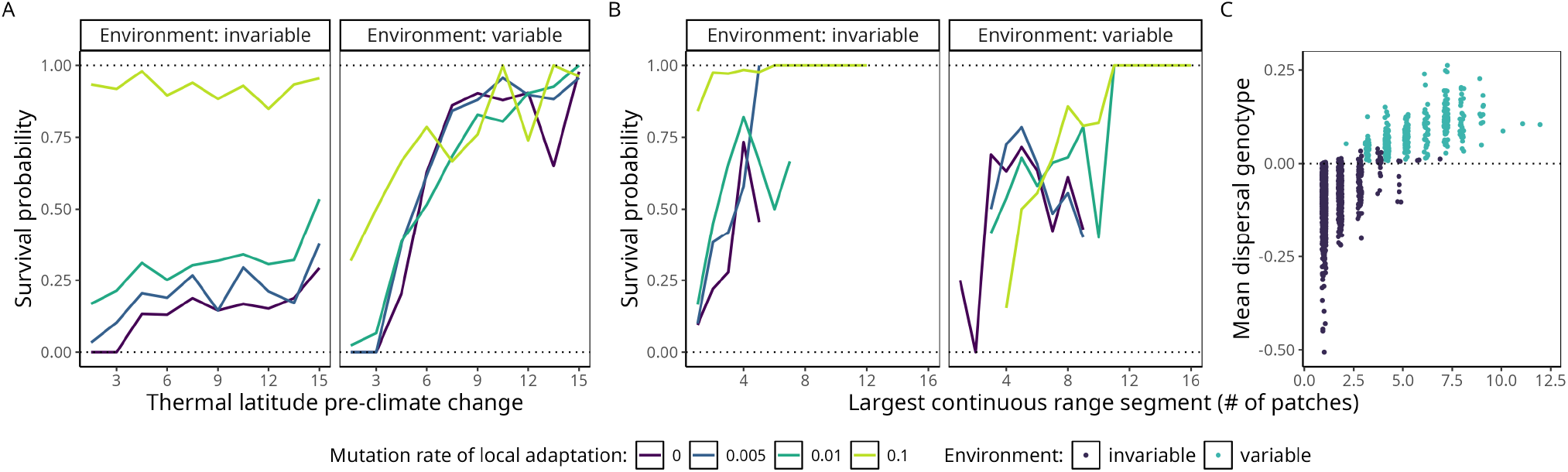
Determinants of species survival under climate change. (A) Survival probability of a species depending on the temperature in its range centroid pre-climate change. (B) Survival probability of a species depending on the size of the largest continuous segment of its range. (C) The genotypic mean of the dispersal trait of a species against its largest continuous range segment just before climate change (t=10,000) with a mutation rate of local adaptation of *m*_*la*_ = 0.005. For all graphs, data comes from 50 replicate simulations. Probabilities are calculated for values of the independent variables with a sample size greater than two. Dispersal mortality is set to *µ*_0_ = 10% and the mutation rate of dispersal during climate change is set to *m*_*d*_ = 0.0005. In invariable environments, environmental stochasticity is set to *σ* = 0.5. In variable environments, environmental stochasticity is set to *σ* = 1.5.

The size and fragmentation of a species’ range pre-climate change is the second important determinant of its survival. Generally, species with smaller and more fragmented ranges are more likely to go extinct during climate change (Figure 4B), matching field observations [48]. In our model, the reason likely lies in the selection pressures on dispersal and the sexual nature of reproduction: Species with isolated ranges could face mate-finding Allee effects during range expansions, which would select further against dispersal [49]. This dynamic mirrors observations of island and mountaintop populations experiencing strong dispersal reduction over time [50]. Once the climate starts changing, such species would neither have the dispersal abilities nor the neighbouring populations to complete successful poleward range shifts. Our results do not depend on changes in dispersal mortality and environmental stochasticity, barring extremely chaotic environments (Figure S9).

The historical environment in which species have evolved is crucial for understanding the survival of species during climate change (Figure 4B,C): Variable environments exclusively produce dispersive species with larger and more connected ranges that can respond relatively well to climate change. In invariable environments, interspecific differences in range isolation and dispersal abilities make survival more conditional on historical range formation and stochastic processes during community assembly (see Figure S10 for an illustration).

More globally, biodiversity post-climate change converges to similar levels almost regardless of the level of environmental stochasticity or evolutionary potential of local adaptation, even though diversity pre-climate change differs wildly between scenarios (Figure 3A). In the absence of complete evolutionary rescue (Figure 3A, green), both the absolute and the relative impact of climate change is strongest for those metacommunities that experienced the least environmental stochasticity pre-climate change. This result is reminiscent of the findings of Bell & Gonzalez [51] and Gonzalez & Bell [52] who show experimentally that evolutionary rescue is most likely to occur in metapopulations with a history of environmental stress. The analogue in our model is that only those species who have evolved to survive high levels of environmental stochasticity also survive climate change.

Overall, our results demonstrate that climate change causes biodiversity loss and forces range expansions. Importantly, we have also shown that the impact of climate change depends heavily on the environmental history and subsequent evolution of dispersal in the metacommunity. The metacommunity’s history will also determine the extent to which contemporary dispersal evolution can rescue species from climate change.

### Dispersal evolution can rescue some, but not all species

Dispersal evolution has some capacity to rescue species during climate change, but it depends on the evolutionary potential of both traits, local adaptation and dispersal. When species can track their niche via local adaptation, increasing the mutation rate of dispersal has no effect (Figure 5A, green lines). When the evolutionary potential for local adaptation is relatively low, biodiversity maintained during climate change increases with the mutation rate of dispersal in invariable environments (Figure 5A, left panel). There is no such effect in variable environments (Figure 5A, right panel). However, the amount of species lost to climate change dwarfs the amount of species rescued by dispersal evolution (Figure 5B, left panel). The results do not depend on changes in dispersal mortality and environmental stochasticity as long as the environment does not become intolerably chaotic (Figure S11). The results are also insensitive to changes in the speed and volume of climate change (Figure S12) but suggest an interaction with local adaptation.

**Figure 5:**
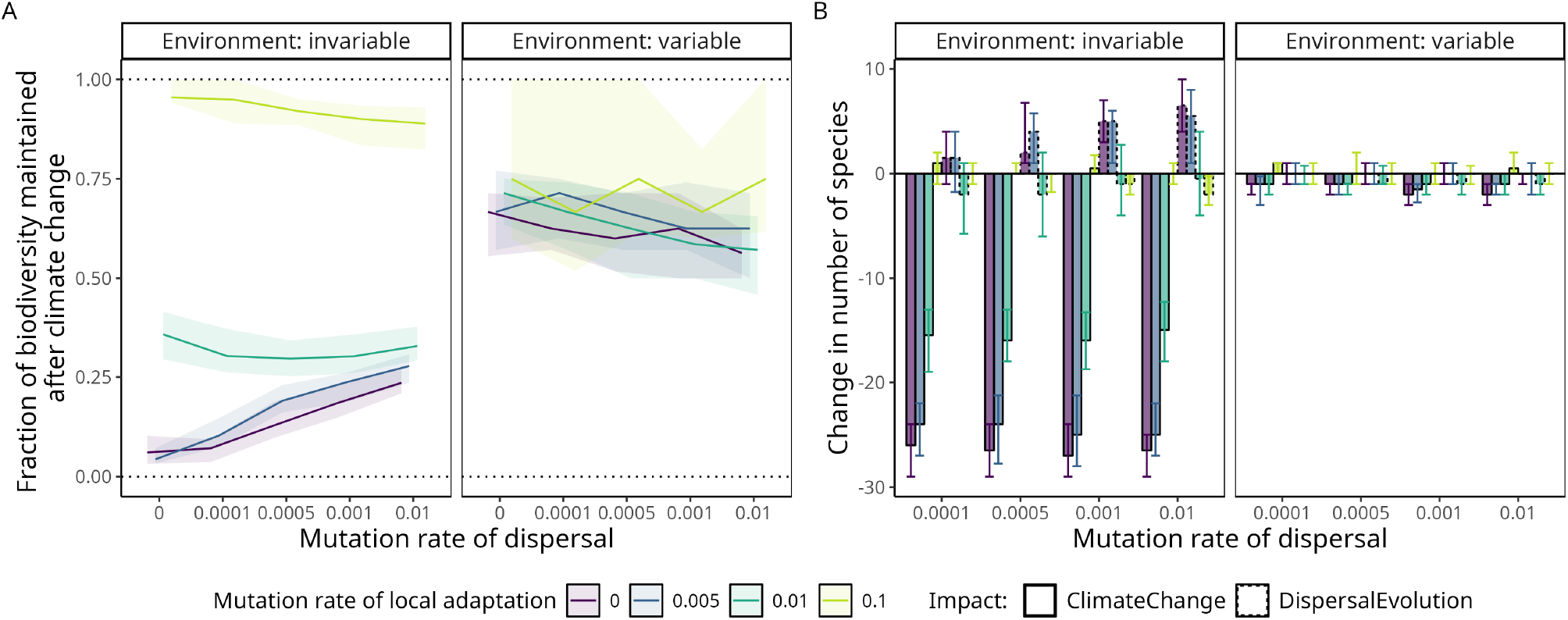
The impact of dispersal evolution on biodiversity maintenance under climate change. Lines/bars represent medians and shaded areas/error bars interquartile ranges across 50 replicate simulations. Dispersal mortality is set to *µ*_0_ = 10%. In invariable environments, environmental stochasticity is set to *σ* = 0.5. In variable environments, environmental stochasticity is set to *σ* = 1.5. (A) The proportion of *γ*-diversity maintained after climate change (t=15,000) compared to before (t=10,000). (B) The change in *γ*-diversity during climate change due to the ecological effects of climate change, compared to the change due to dispersal evolution. We disentangle the species lost due to climate change and the species rescued due to dispersal evolution using a benchmark scenario, in which we do not allow for dispersal evolution during climate change (see Figure S3).

Community rescue in invariable environments depends on high mutation rates (Figure 5A, left panel) because robust dispersal traits emerge in these environments. A trait is robust when genotypic changes due to mutation or recombination do not incur significant phenotypic changes [35, 53]. Whether a trait can evolve robustness depends on the genotype-to-phenotype map [54]. In our model, we assume that the dispersal trait is not bounded at the genotypic level. However at the phenotypic level, the dispersal trait — a probability — necessarily takes values between 0 and 1. This genotype-phenotype map implies that, under conditions that select very strongly and consistently against dispersal (such as invariable environments, Figure 2B), robustness can emerge: The genotype can take values below zero, implying that a phenotypic change away from zero is more difficult (Figure 4C). In our model, robustness occurs only in those species that have faced very strong selection pressures against dispersal in their evolutionary history (see Figure S10 for a comparison of genotypic trajectories between invariable and variable environments). This is representative of cases in which robustness should occur in natural populations, as it has been shown to evolve rather rapidly in the presence of strong selection pressures [55].

Previous work on robustness and evolvability has demonstrated that the genotype-to-phenotype map and its consequences for mutational effects can have eco-evolutionary implications up to the landscape scale [56, 57]. While lacking in sophistication, our genotype-to-phenotype map gives us the intuition that certain environmental and range characteristics might select so strongly against dispersal that it becomes difficult to regain these abilities once lost. In natural populations, for example, there is broad empirical support for loss of flight ability in many insects and birds that are isolated on islands or in habitat fragments [50]. Re-evolution of wings on the other hand is very rare — which may suggest robustness, but this remains of course speculative [50, 58]. Similarly, there is evidence for reduced and robust dispersal abilities at higher elevations, co-occurring with isolated populations in mountaintop habitats [59]. Our results suggest that such robustness of dispersal due to species’ environmental and evolutionary histories could severely constrain their potential of rescue via dispersal evolution during climate change.

More generally, rescue effects via dispersal evolution have been shown in other theoretical work [22, 60], as has the rapid evolution of dispersal at range edges in natural populations [27, 61]. The evidence for the capacity of natural populations to track climate change with their niches on the other hand is ambiguous [26, 62], but such evolutionary rescue via local adaptation has been documented in freshwater fish [63], *Drosophila* flies [64], and marine phytoplankton [65]. Depending on species’ life- and environmental histories, rescue via contemporary dispersal evolution is likely a salient but limited factor in predicting biodiversity loss due to climate change.

### When is contemporary dispersal evolution relevant?

The ultimate goal of this article is to delineate the space in which contemporary dispersal evolution is relevant for biodiversity maintenance under climate change. To be relevant, it is necessary that rapid dispersal evolution enables range shifts that would not have been possible otherwise. In the absence of complete evolutionary rescue via local adaptation, there are thus three general cases to distinguish: (i) Range shifts are possible with the historically evolved dispersal abilities, (ii) range shifts are impossible, and (iii) range shifts become possible with rapid dispersal evolution. Figure 6 conceptualises these three cases and their eco-evolutionary origins.

**Figure 6:**
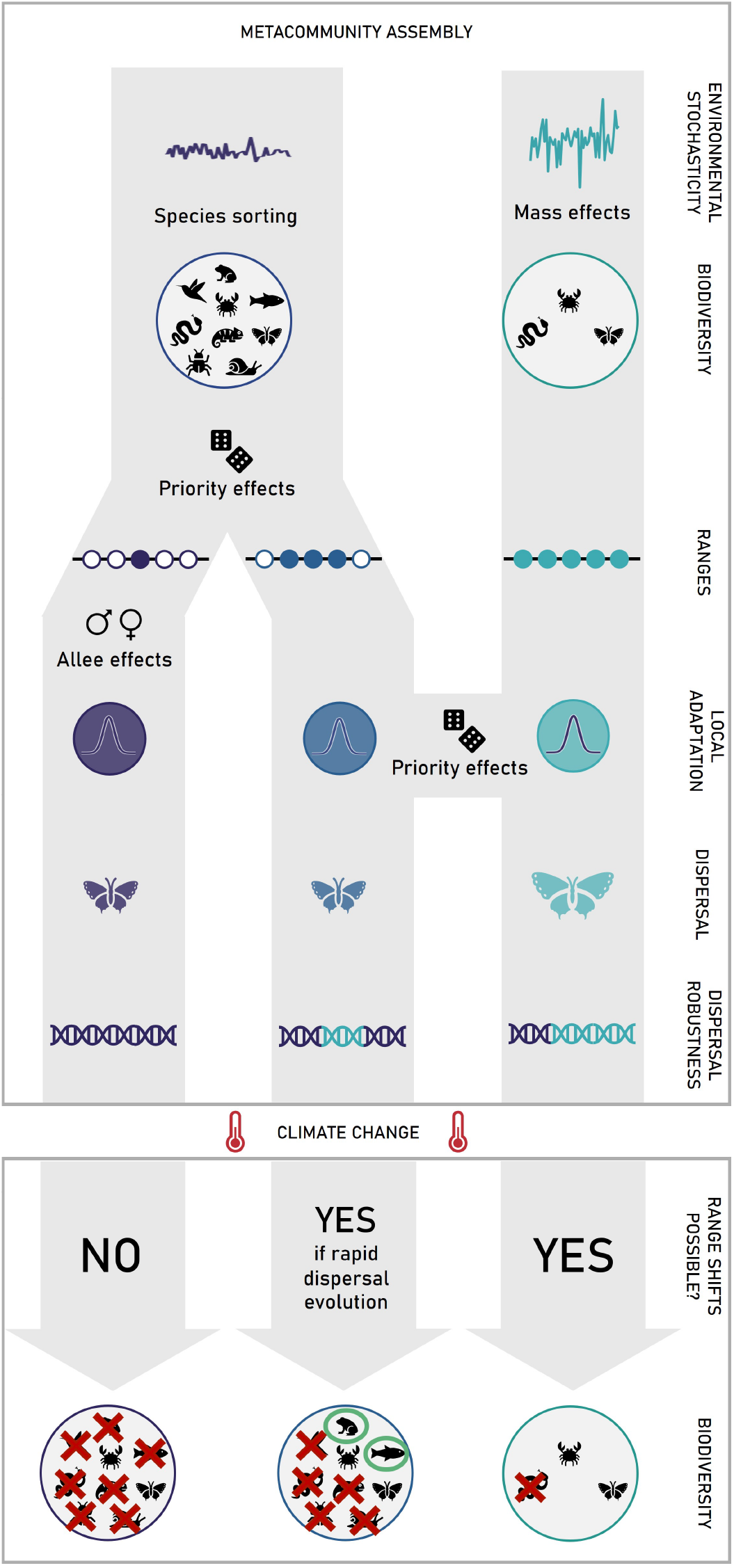
Conceptual framework of eco-evolutionary metacommunity dynamics during assembly from the landscape to the gene level, and the subsequent implications under climate change. At the bottom, red crosses signify species extinctions, green circles signify species rescue via rapid dispersal evolution.

The first case (range shifts are possible with historically evolved dispersal abilities, right side of Figure 6) appears mostly in historically variable environments. Community assembly is driven by spatial bet-hedging [19, 39] and species evolve relatively high dispersal rates and large ranges. Thus, mass effects occur during assembly [17, 42, 43], and communities are quite similar across space and not diverse overall. In this case, range shifts are likely possible with the evolved dispersal abilities. If any standing genetic variation is left in the population, those range shifts could be sped up via rapid evolution. However, our model predicts that the evolved dispersal abilities should be sufficient for species to track their environmental niche in space and biodiversity losses due to climate change should be rather small. Typical examples of this case could be metacommunities of northern European butterflies: They are characterised by relatively strong extinction-colonization dynamics [66], and climate change-induced range expansions have been shown in multiple species [27].

The second case (range shifts are impossible, left side of Figure 6) is the most catastrophic for biodiversity. It most likely occurs in historically invariable environments, in which dispersal is selected against [39] and species sorting [17, 42, 43] occurs. Once climate change starts, species do not have the necessary dispersal abilities to shift their ranges, and even if some individuals were to disperse, they would have to expand their range facing mate-finding Allee effects — if reproduction is sexual as we assume here. This case most likely occurs in isolated species with concordant ancient and strong selection that has reduced genetic variation to a minimum. Even if mutations could occur quickly enough, this stabilizing selection would likely produce dispersal traits that are canalised, that is, robust to mutations [35]. Natural examples include island and mountaintop populations, characterised by high degrees of endemism and isolation over long periods of time [67, 68], but see [69].

The third case (range shifts are possible with rapid evolution of dispersal, centre of Figure 6) is more nuanced. In relatively invariable environments with species sorting dynamics, stochastic and historical interspecific differences can produce variable range sizes (Figure S10). For the few species that occupy comparatively large ranges in such environments, it depends on their historical presence in the range and subsequent local adaptation. Species with ancient and well-adapted local populations experience strong selection against dispersal, while species with newer and less well-adapted populations are still dispersive across the range (Figure S13). The latter scenario resembles the first case, depicted on the right side of Figure 6: Range shifts are possible with the evolved dispersal abilities. The former scenario captures the distinct subset of species that can theoretically benefit from contemporary dispersal evolution during climate change: Species from invariable environments, with large ranges and well-adapted local populations. If dispersal evolution is rapid enough, individuals from these species can disperse into neighbouring populations and thus shift their range. Examples of natural systems for this case may include species with strongly peaked fitness surfaces across their geographical range, as recently shown in *Drosophila* [70], but this remains speculative.

It is important to note that in our model, the capacity of the system to respond to climate change mainly depends on new variation introduced by mutations since existing variation is usually limited after community assembly. Some theoretical work predicts the rapid evolution of increased mutation rates during range expansions [56], but such *de novo* mutations of dispersal may be too limited in their effect to fundamentally change a population’s response on ecological timescales [50]. Rapid evolution of dispersal during range expansions, as documented in natural and experimental systems, likely acts on standing genetic variation [61, 71]. In natural systems, such genetic variation can be maintained through other environmental and genetic mechanisms such as dominance or epistasis [72, 73]. It is possible that selection on such genetic variation can lead to rescue effects via dispersal evolution in natural populations, and the same holds for phenotypic plasticity [74] or epigenetic mechanisms [75]. However, our work highlights that information on genotype-phenotype maps and trait robustness is a necessary complement to information on standing genetic variation and mutation rates if we want to predict how biodiversity responds to climate change. The lack of empirical data in this context is worrying and our work underlines the importance of such integration across scales, from genomes to species ranges.

Distinguishing the three cases above maps out a clear framework for understanding community responses to climate change: Knowledge of the environmental and evolutionary history of a given community is crucial, as is an understanding of the existing evolutionary potential, or conversely, the evolved robustness. These results stress the empirically supported importance of evolutionary legacy effects for responses to contemporary change [76]: For example, local adaptation has been shown to depend on environmental history [51, 52], and thermal performance evolution has been shown to depend on the community background [77]. We elucidate here a mechanism by which such evolutionary legacy effects can effectively constrain the potential for dispersal evolution to rescue species from climate change.

To take into account the effects of species’ evolutionary legacies, efforts to predict the capacity for climate change-induced range shifts should not just measure current dispersal abilities, but also the variation in the traits determining these abilities, species isolation, and extinction-colonization dynamics. Such a mechanistic understanding of the capacity for range shifts is particularly important for species in which the potential for sufficiently rapid climatic niche evolution remains questionable [26, 28–30].

## Conclusion

Anthropogenic climate change is accelerating, and so are its impacts on global biodiversity [78]. Here, we show that dispersal evolution and its drivers are key factors in understanding both the history of ecological communities as well as their capacity to respond to climate change. Historically variable environments equip species with sufficiently large ranges and dispersal abilities for range shifts towards colder climates. Species from historically invariable environments depend on rapid evolution of dispersal for their survival during climate change. However, their rescue might be constrained by the evolution of robust dispersal traits. Conservation efforts such as assisted gene flow [79] may therefore be most necessary for species with smaller, more fragmented, or colder ranges with a history of relatively stable environments.

## Data Availability

Computer code is available at GitHub and Zenodo (DOI: https://doi.org/10.5281/zenodo.15228385).

## Author contributions

E.A.F., P.L.T and P.K. designed the research. P.K., N.L. and E.A.F. performed research. P.K. and N.L. analyzed data. P.K. wrote the paper and all authors commented on the manuscript.

## Acknowledgments

We thank Juliane Mailly and Jhelam Deshpande for helpful discussions. This work was supported by grants from the Agence Nationale de la Recherche (Grants ANR-19-CE02-0015 and ANR-23-CE02-0030).

## Notes

### Competing Interest Statement

The authors have declared no competing interest.

### Summary of Updates

Updated manuscript for more clarity and accessibility, added Figure 1.

