## Supplementary Material for "Dispersal evolution can only rescue a limited set of species from climate change"

### Supplementary Information for: Dispersal evolution can only rescue a limited set of species from climate change

April 16, 2025

#### Supplementary Tables (Table S1)

Table S1: Explored parameters and their ranges of values

| Parameter | Description | Explored (standard) values |
|---|---|---|
| $\mu_0$ | Dispersal mortality | 0,0.001,0.01,0.05,0.1,0.2,0.3,0.4,0.5 |
| $\sigma$ | Environmental stochasticity | 0,0.25,0.5,0.75,1,1.25,1.5,1.75,2 |
| $\epsilon$ | Patch extinction probability | 0,0.001,0.003,0.005,0.01,0.02,0.03,0.04,0.05 |
| $m_d$ | Mutation rate of dispersal (during climate change) | 0,0.0001,0.0005,0.001,0.01 |
| $m_{la}$ | Mutation rate of local adaptation | 0,0.005,0.01,0.1 |
| $\delta_{envchange}$ | Volume of climate change per generation | 0.0005,0.001,0.002 |

### Supplementary Figures

#### Methods (Figures S1-S3)

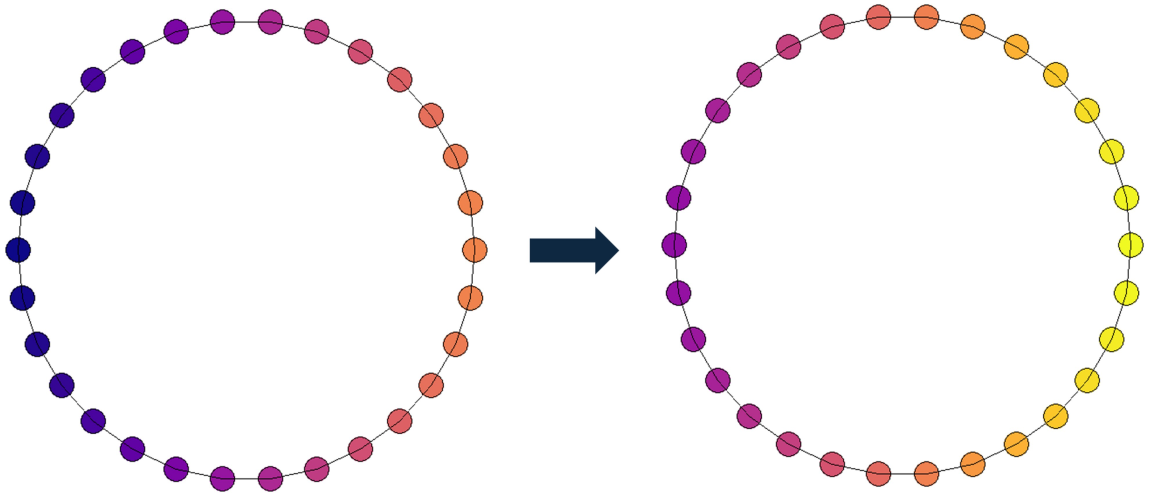

Figure S1: Graphical representation of the landscape: A ring of patches with a temperature gradient. Temperatures increase uniformly during climate change (from left to right).

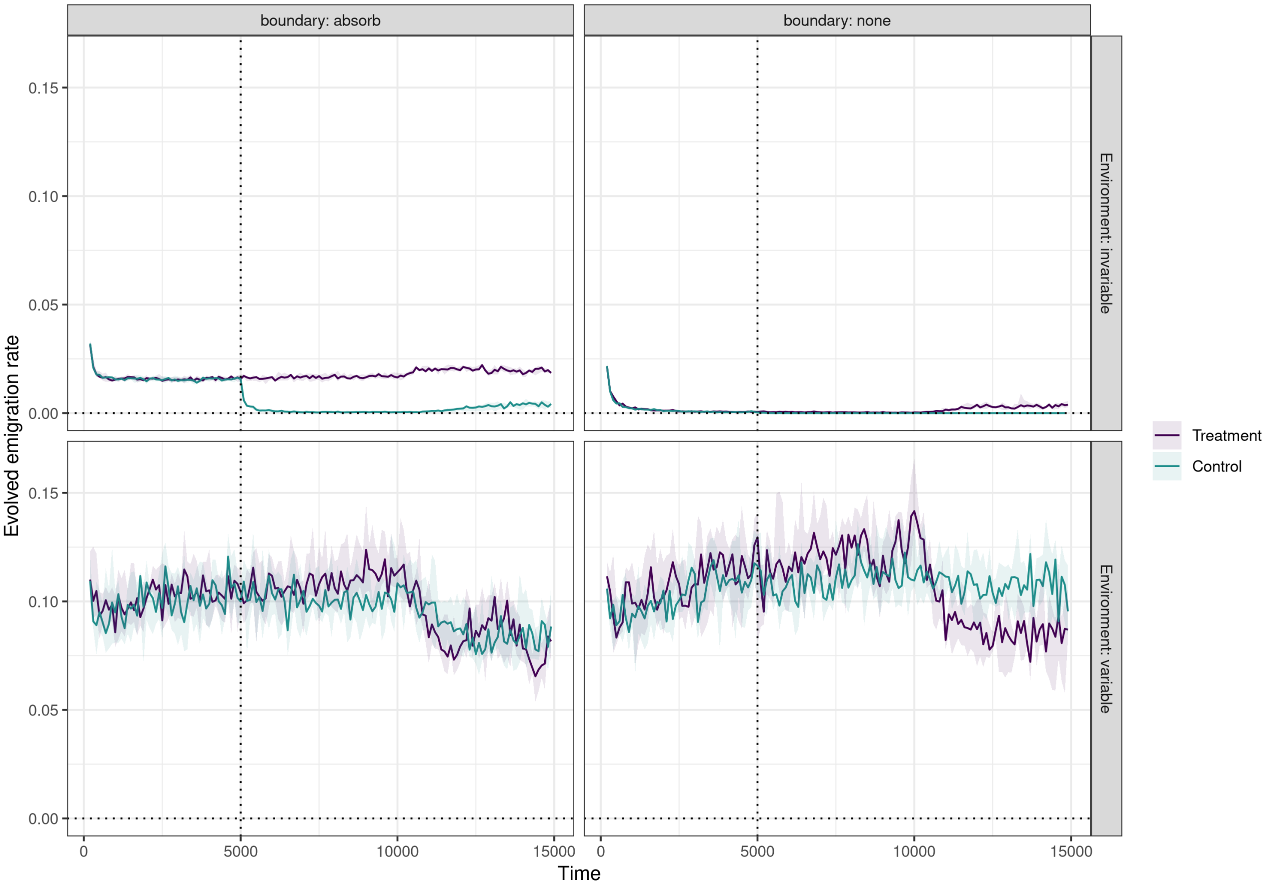


Figure S2: Comparison of two boundary conditions of the dispersal genotype. The left column is with absorbing boundaries i.e., when the trait crosses the 0 or 1 it gets set to 0 or 1, respectively. The right column is with no boundaries. In invariable environments, environmental stochasticity is set to $\sigma = 0.5$. In variable environments, environmental stochasticity is set to $\sigma = 1.5$. The outcome variable is the emigration rate in the metacommunity over time. Lines represent medians and shaded areas interquartile ranges across 10 replicate simulations. Treatment (purple) describes a case in which dispersal mutates with a rate of $m_d = 0.01$ all throughout the simulation. Control (green) describes a case in which mutations for dispersal stop after time step 5,000 (see dotted line). In invariable environments, there is a clear quantitative difference in the stable dispersal rate between boundary conditions. Additionally, when we stop mutations at time step 5,000 in the control scenario, the emigration rate of the absorbing boundary drops to the level of the no boundary case, where there are no large differences between treatment and control. This implies that the mutation-selection balance biases our dispersal rates upwards if we introduce absorbing boundaries. This is much more pronounced for dispersal rates closer to 0 (compare to the variable environment), and therefore likely an artefact of the genetic algorithm trying to avoid the 0 in the absorbing boundary case, where it constitutes a mathematical attractor.

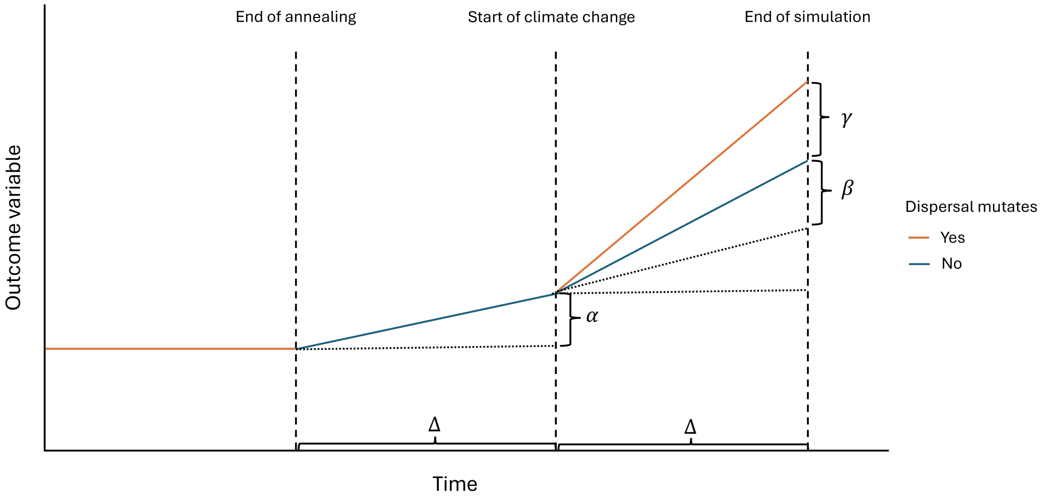


Figure S3: Illustration of the decomposition of the observed impacts. $\alpha$ serves as the estimate of the impact of the environment absent dispersal evolution or climate change, $\beta$ as the impact of climate change without dispersal evolution or environmental factors, and $\gamma$ as the impact of dispersal evolution.

#### Community assembly (Figures S4-S6)

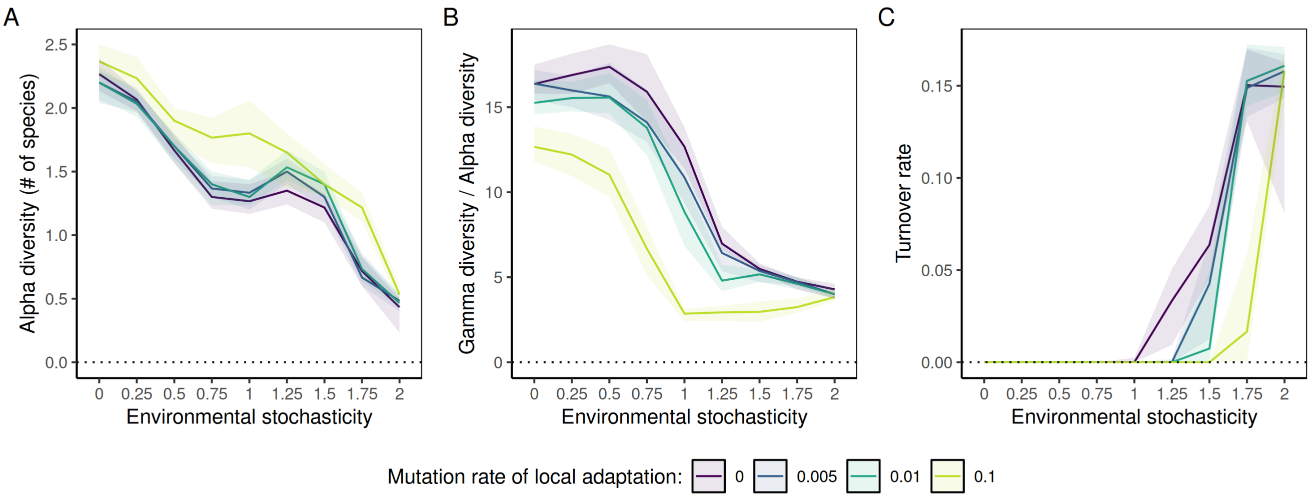


Figure S4: Additional outcomes of community assembly dependent on environmental stochasticity. Lines represent medians and shaded areas interquartile ranges across 50 replicate simulations at the end of community assembly (t=10,000). Dispersal mortality is set to an intermediate $\mu_0 = 10\%$. Outcome variables are: (A) $\alpha$-diversity i.e., mean number of species per patch; (B) $\gamma$-diversity divided by $\alpha$-diversity, high values indicate community dissimilarity; (C) the average turnover rate during the last 100 generations of community assembly, calculated as the ratio of patches that change occupancy status in one time step.

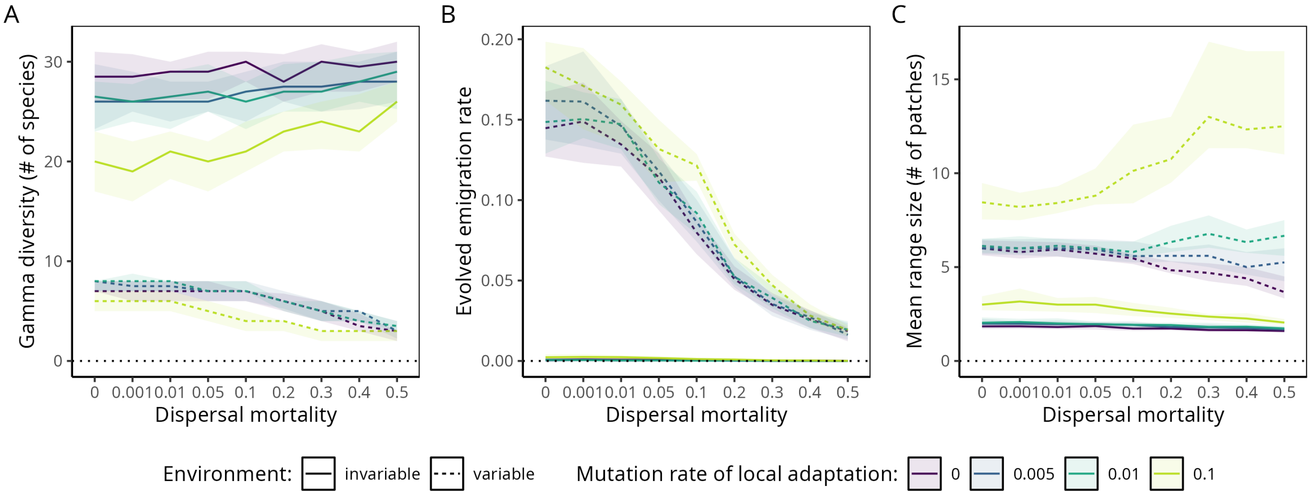


Figure S5: Community assembly dependent on dispersal mortality $\mu_0$. Lines represent medians and shaded areas interquartile ranges across 50 replicate simulations at the end of community assembly (t=10,000). In invariable environments, environmental stochasticity is set to $\sigma = 0.5$. In variable environments, environmental stochasticity is set to $\sigma = 1.5$. Outcome variables are: (A) $\gamma$-diversity i.e., total number of species in the metacommunity; (B) the average emigration rate in the metacommunity; (C) the average range size of a species in number of patches.

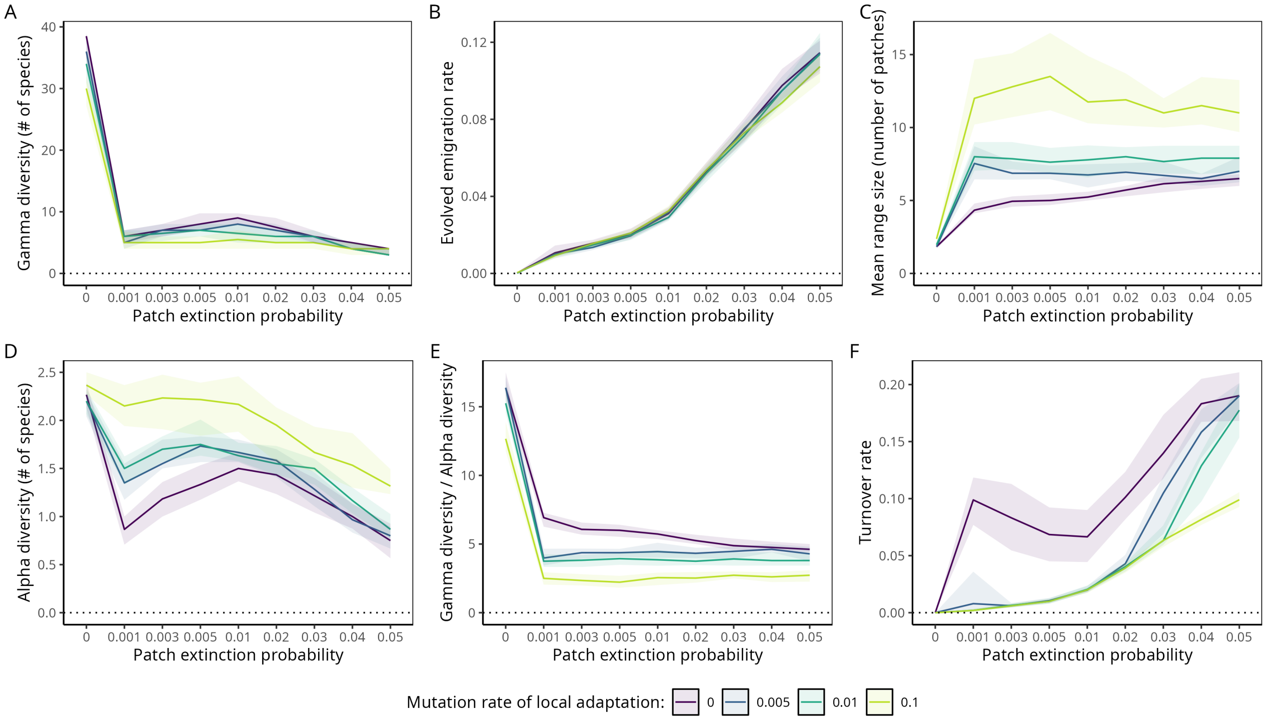


Figure S6: Alternative results for community assembly utilizing random patch extinctions to generate environmental variation. Lines represent medians and shaded areas interquartile ranges across 50 replicate simulations at the end of community assembly (t=10,000). Dispersal mortality is set to an intermediate $\mu_0 = 10\%$. Outcome variables are: (A) $\gamma$-diversity i.e., total number of species in the metacommunity; (B) the average emigration rate in the metacommunity; (C) the average range size of a species in number of patches; (D) $\alpha$-diversity i.e., mean number of species per patch; (E) $\gamma$-diversity divided by $\alpha$-diversity, high values indicate community dissimilarity; (F) the average turnover rate during the last 100 generations of community assembly, calculated as the ratio of patches that change occupancy status in one time step.

#### Impact of climate change (Figures S7-S8)

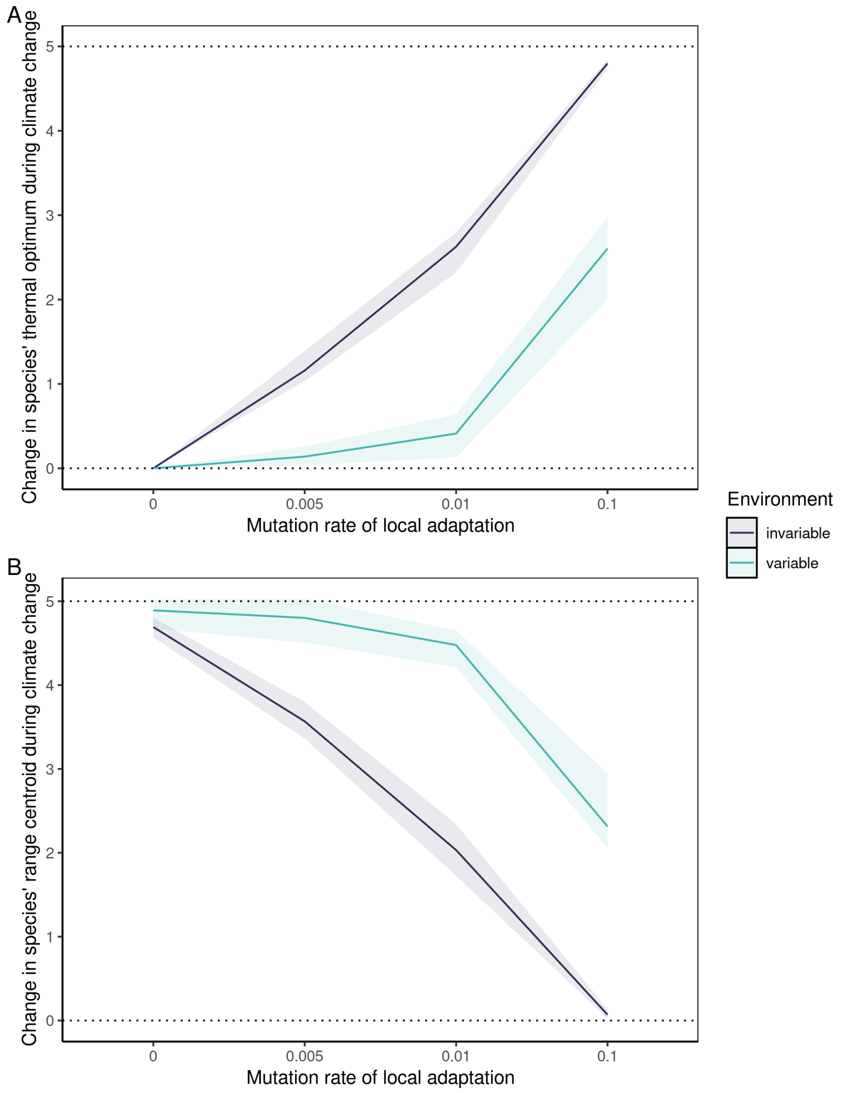


Figure S7: Niche and range shifts during climate change depending on the potential for local adaptation and the amount of environmental stochasticity. Lines represent medians and shaded areas interquartile ranges across 50 replicate simulations. Outcomes are simple differences between time steps t=10,000 and t=15,000. The total volume of environmental (climate) change is 5 (see dotted lines). The mutation rate of dispersal is set to $m_d = 0.0005$, the dispersal mortality is set to $\mu_0 = 10\%$. In invariable environments, environmental stochasticity is set to $\sigma = 0.5$. In variable environments, environmental stochasticity is set to $\sigma = 1.5$. (A) describes the average shift in a species' thermal optimum i.e., in the trait coding for local adaptation. (B) describes the average shift of a species' range centroid. Positive values indicate shifts towards cooler climates.

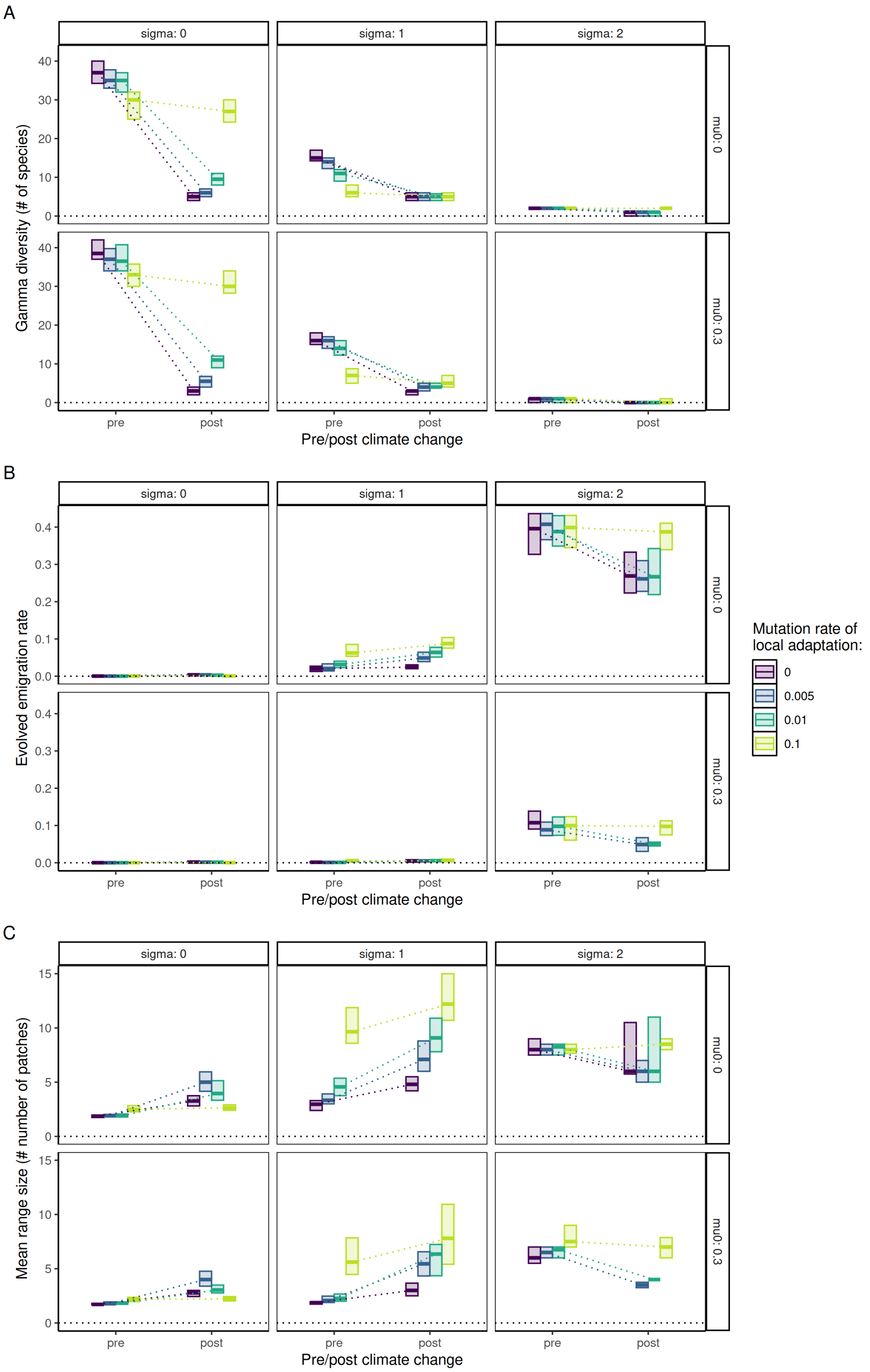


Figure S8: Sensitivity analysis for the impact of climate change, varying environmental stochasticity (sigma) and dispersal mortality (mu0) beyond the standard scenarios presented in the main text. Thick lines represent medians and shaded boxes interquartile ranges across 50 replicate simulations. The mutation rate of dispersal is set to $m_d = 0.0005$. (A) $\gamma$-diversity i.e., total number of species in the metacommunity; (B) the average emigration rate in the metacommunity; (C) the average range size of a species in number of patches.

#### Determinants of survival (Figure S9)

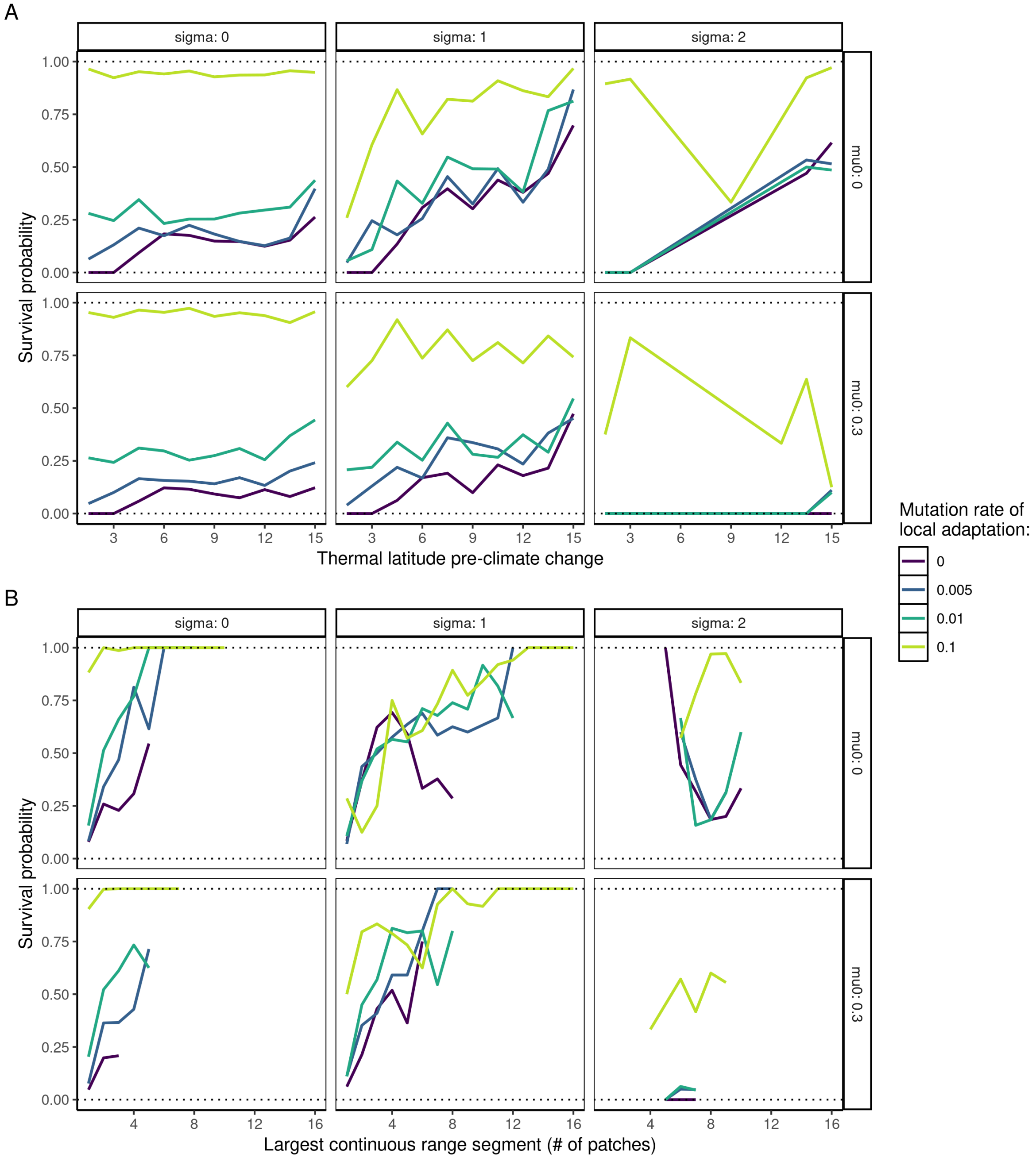


Figure S9: Sensitivity analysis for the determinants of species' survival, varying environmental stochasticity (sigma) and dispersal mortality (mu0) beyond the standard scenarios presented in the main text. All data points are probabilities across 50 replicate simulations. Probabilities are calculated for values of the independent variables with a sample size greater than two. The mutation rate of dispersal is set to $m_d = 0.0005$. (A) Survival probability of a species depending on the temperature in its range centroid pre-climate change. (B) Survival probability of a species depending on the size of the largest continuous segment of its range.

#### Impact of dispersal evolution (Figure S10-S13)

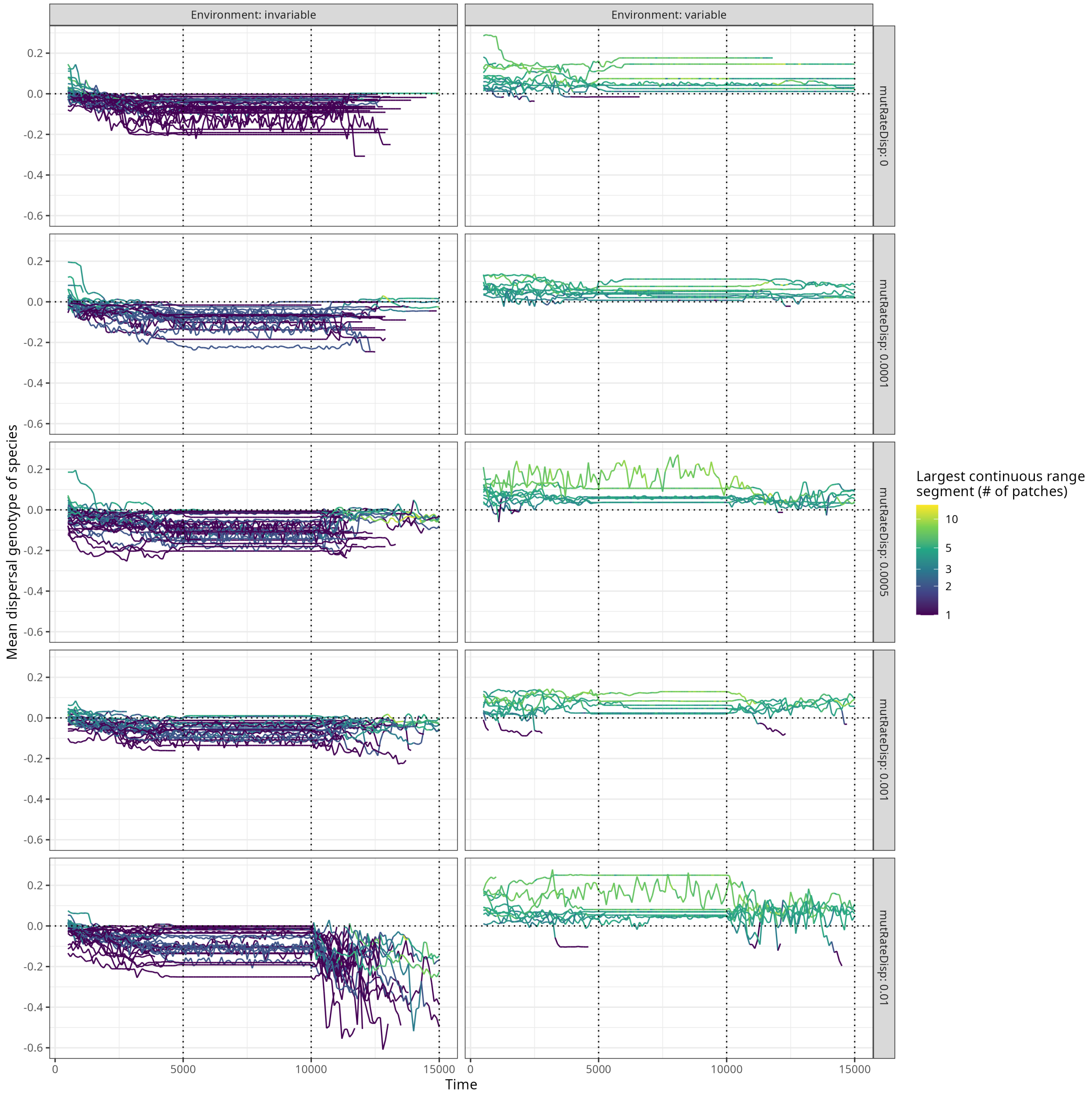


Figure S10: Mean dispersal genotype per species over time. Data is from one representative replicate run. The mutation rate of local adaptation is set to $m_{la} = 0.005$, the dispersal mortality is set to $\mu_0 = 10\%$. In invariable environments, environmental stochasticity is set to $\sigma = 0.5$. In variable environments, environmental stochasticity is set to $\sigma = 1.5$. The mutation rate of dispersal varies on the global y-axis. Each line represents one species, the color of the line represents the size of the largest continuous segment of this species range. The dotted vertical lines represented important markers: the end of annealing (t=5,000), the start of climate change (t=10,000), and the end of the simulation (t=15,000). The negative trend in the bottom left panel stems from an increased mutation rate compared to the assembly phase and represents an increase of the trait variance in the space below zero.

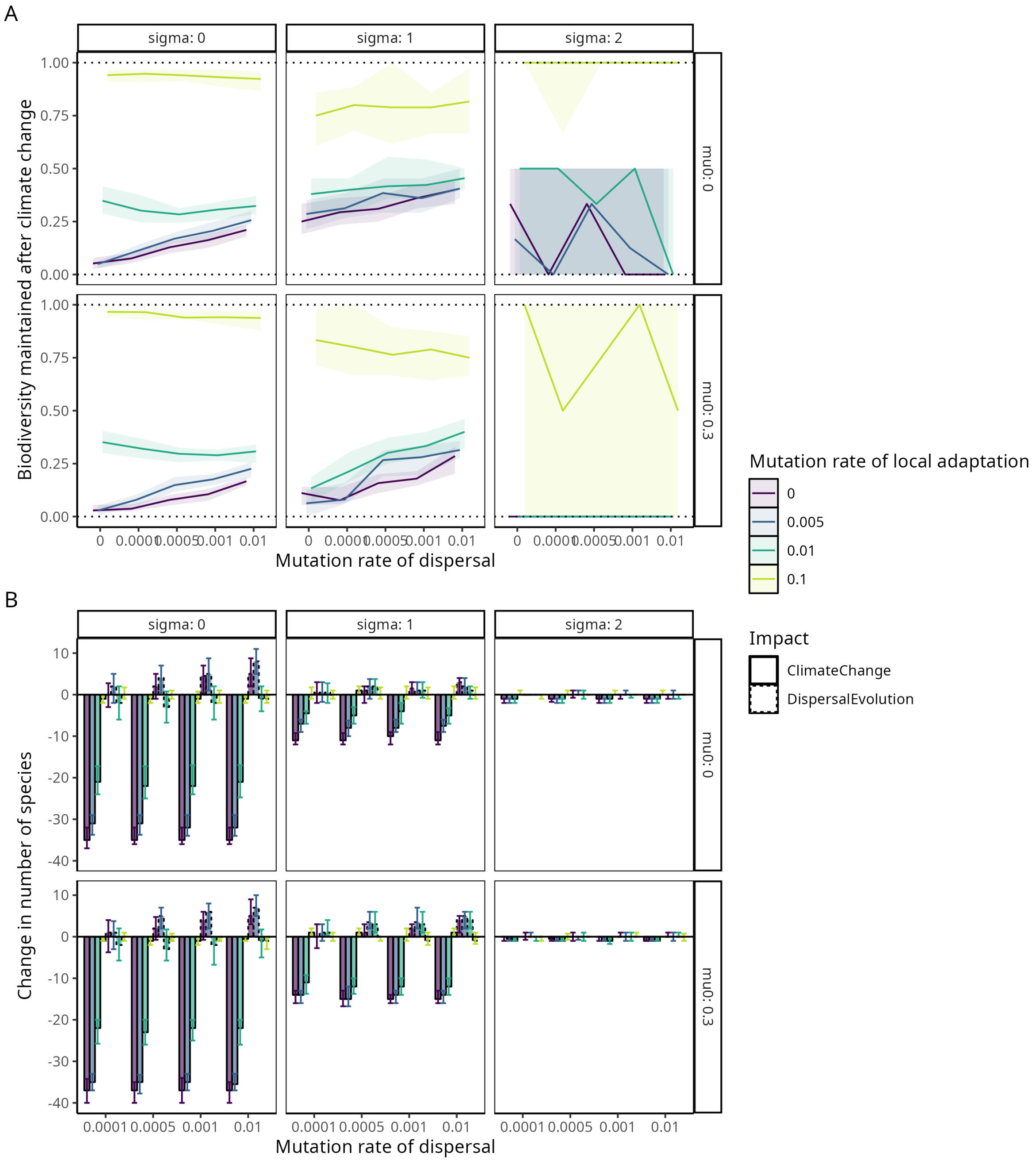


Figure S11: Sensitivity analysis for the impact of dispersal evolution under climate change, varying environmental stochasticity (sigma) and dispersal mortality (mu0) beyond the standard scenarios presented in the main text. Lines/bars represent medians and shaded areas/error bars interquartile ranges across 50 replicate simulations. (A) The proportion of $\gamma$-diversity maintained after climate change (t=15,000) compared to before (t=10,000). (B) The change in $\gamma$-diversity during climate change due to the ecological effects of climate change, compared to the change due to dispersal evolution. The effect of climate change is obtained from a benchmark scenario, in which the mutation rate of dispersal is set to $m_d = 0$ (see Supplementary Methods).

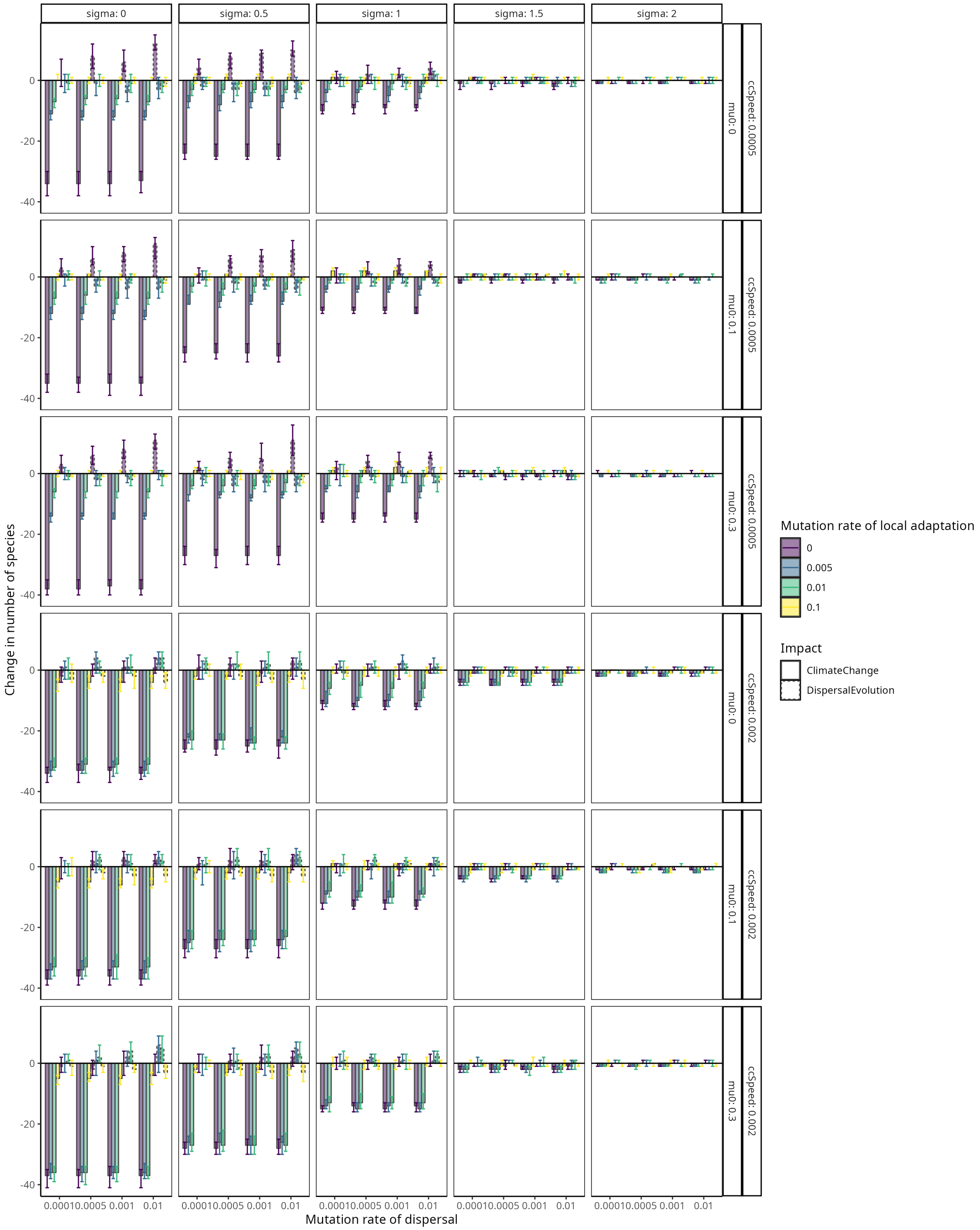


Figure S12: Sensitivity analysis for the impact of dispersal evolution under climate change, varying the speed of climate change (ccSpeed), environmental stochasticity (sigma), and dispersal mortality (mu0) beyond the standard scenarios presented in the main text. Bars represent medians and error bars interquartile ranges across 50 replicate simulations. (A) The proportion of $\gamma$-diversity maintained after climate change (t=15,000) compared to before (t=10,000). (B) The change in $\gamma$-diversity during climate change due to the ecological effects of climate change, compared to the change due to dispersal evolution. The effect of climate change is obtained from a benchmark scenario, in which the mutation rate of dispersal is set to $m_d = 0$ (see Supplementary Methods).

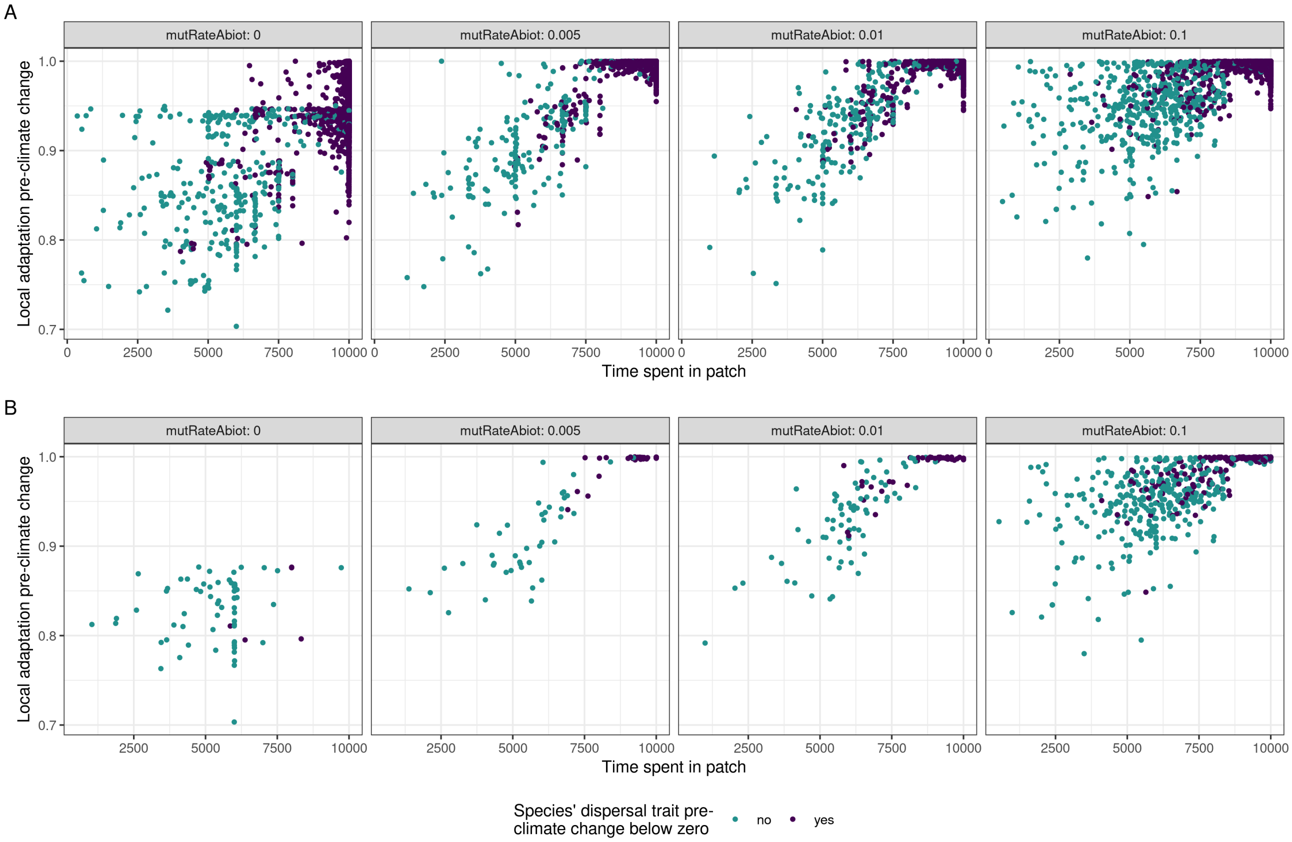


Figure S13: Local adaptation of a species pre-climate change (t=10,000, see Supplementary Methods for calculation of local adaptation) against the average time the species has inhabited all of the patches in its range at t=10,000. (A) Results for all species, (B) Results only for species with ranges greater than 5 patches. The mutation rate of local adaptation is varied in the facets. The color scale indicates whether local adaptation has selected so strongly against dispersal that the species' average dispersal trait has drifted below zero. Data are from 50 replicate simulations in invariable environments (environmental stochasticity is set to $\sigma = 0.5$, dispersal mortality to $\mu_0 = 10\%$).
